# Physachenolide C induces complete regression of established murine melanoma tumors via apoptosis and cell cycle arrest

**DOI:** 10.1101/2021.05.27.445710

**Authors:** Anngela C. Adams, Anne M. Macy, Paul Kang, Karla F. Castro-Ochoa, E. M. Kithsiri Wijeratne, Ya-Ming Xu, Manping X. Liu, Alexandra Charos, Marcus W. Bosenberg, A. A. Leslie Gunatilaka, Aparna R. Sertil, K. Taraszka Hastings

## Abstract

Melanoma is an aggressive skin cancer that metastasizes to other organs. While immune checkpoint blockade with anti-PD-1 has transformed the treatment of advanced melanoma, many melanoma patients fail to respond to anti-PD-1 therapy or develop acquired resistance. Thus, effective treatment of melanoma still represents an unmet clinical need. Our prior studies support the anti-cancer activity of the 17β-hydroxywithanolide class of natural products, including physachenolide C (PCC). As single agents, PCC and its semi-synthetic analog demonstrated direct cytotoxicity in a panel of murine melanoma cell lines, which share common driver mutations with human melanoma; the IC_50_ values ranged from 0.18 – 1.7 µM. PCC treatment induced apoptosis of tumor cells both *in vitro* and *in vivo. In vivo* treatment with PCC alone caused the complete regression of established melanoma tumors in all mice, with a durable response in 33% of mice after discontinuation of treatment. T cell-mediated immunity did not contribute to the therapeutic efficacy of PCC or prevent tumor recurrence in YUMM2.1 melanoma model. In addition to apoptosis, PCC treatment induced G0-G1 cell cycle arrest of melanoma cells, which upon removal of PCC, re-entered the cell cycle. PCC-induced cycle cell arrest likely contributed to the *in vivo* tumor recurrence in a portion of mice after discontinuation of treatment. Thus, 17β-hydroxywithanolides have the potential to improve the therapeutic outcome for patients with advanced melanoma.

## Introduction

Melanoma is an aggressive skin cancer with increased risk of metastasis if it is not treated at an early stage. Immune checkpoint blockade has transformed the treatment of advanced melanoma, defined as unresectable or metastatic disease. However, in the first line treatment setting, only 33-44% of patients respond to monoclonal antibodies (mAb) targeting PD-1 (Larkin, et al., 2015, Robert, Long, et al., 2015, Robert, Schachter, et al., 2015) and 58% of patients respond to combination therapy with anti-PD-1 and anti-CTLA-4 (Larkin, et al., 2015). Additionally, 25-40% of initial responders develop acquired resistance to anti-PD-1 (Ribas, et al., 2016, Topalian, et al., 2014, Wang, Eroglu, et al., 2017). Therefore, innovative therapeutic approaches are required for improved patient outcomes in melanoma.

Natural products are an important source of anti-cancer drugs, including well-known and effective chemotherapeutic agents such as paclitaxel, vincristine, adriamycin and vinblastine (Newman and Cragg, 2020). Withanolides are a group of naturally occurring steroidal lactones isolated from some members of the *Solanaceae* plant family (Chen, et al., 2011). This class of natural products has shown promising anti-cancer properties against lung, breast, prostate, pancreatic, ovarian, renal and melanoma cancer cells (Chang, et al., 2007, Kakar, et al., 2017, Mayola, et al., 2011, Tewary, et al., 2021, Wang, et al., 2012, Xu, et al., 2017, Xu, et al., 2015, Yu, et al., 2010). Structural optimization of withanolides has yielded superior drug candidates. For instance, structure-activity relationship studies of 17β-hydroxywithanolides have led to the identification of physachenolide C (PCC) as a more and potent and selective anti-cancer withanolide compared to withaferin A and withanolide E (Xu, et al., 2017). The anti-cancer activity of PCC has been demonstrated in androgen-dependent and -independent prostate cancer mouse models, as well as its cytotoxic activity, with IC_50_ values in the nanomolar range, against multiple human prostate cancer cells, but not to normal foreskin fibroblasts (Xu, et al., 2017, Xu, et al., 2015). The therapeutic effect of PCC in combination with poly I:C has been evaluated in a xenograft model of M14 human melanoma and in the non-immunogenic B16-F10 mouse melanoma model. In both cases, treatment with the combination of PCC and poly I:C resulted in enhanced tumor regression compared to single agents or control (Tewary, et al., 2021). Additionally, PCC enhanced melanoma cell death in response to soluble mediators produced by activated T cells, including TNF-α (Tewary, et al., 2021), suggesting that PCC may improve the therapeutic efficacy of T cell-mediated immunotherapeutic approaches.

Here, we report the *in vitro* cytotoxicity of PCC and its semi-synthetic analogue LG-134, as well as the *in vivo* therapeutic efficacy of PCC in treating established tumors, using clinically-relevant, immunogenic murine melanoma models. Furthermore, we investigate the T cell-mediated contribution to the therapeutic efficacy of PCC and whether T cell-mediated immunotherapy using anti-PD-1 mAb enhances the efficacy of PCC.

## Materials and methods

### Mice

C57BL/6J wild-type and RAG1^-/-^ mice were purchased from Jackson Laboratory (Bar Harbor, ME) or bred for *in vivo* experiments. Mice were maintained in microisolator cages in the vivarium facility at the University of Arizona College of Medicine – Phoenix in accordance with local and national guidelines. All studies were approved by the University of Arizona’s Institutional Animal Care and Use Committee.

### Cell lines

Yale University Mouse Melanoma (YUMM) cell lines YUMM2.1 and YUMMER1.7 were previously generated (Meeth, et al., 2016, Wang, Perry, et al., 2017). Similar to YUMMER1.7, YUMM Exposed to Radiation (YUMMER).G was created by exposing YUMM1.G to ultraviolet light (Meeth, et al., 2016, Ramseier, et al., 2019, Wang, Perry, et al., 2017). Both YUMMER lines mimic the high ultra-violet light-induced mutational burden found in human disease. The parental cell lines were derived from genetically engineered mouse models expressing driver mutations commonly found in human melanoma. YUMM2.1 express Braf^V600E/WT^, Pten^-/-^, Cdkn2a^+/-^, and Bcat^STA/WT^. YUMMER1.7 and YUMMER.G both express BrafV600E/WT, Pten^-/-^, Cdkn2a^-/-^, and YUMMER.G additionally express Mc1r^e/e^. Cells were maintained in DMEM:F12 media supplemented with 10% fetal bovine serum, 1% non-essential amino acids, and 1% penicillin-streptomycin. Cells were tested at the start and conclusion of experiments for mycoplasma contamination by PCR.

### Production of PCC and LG-134

The natural product, physachenolide C (PCC), was obtained by epoxidation of physachenolide D isolated from *Physalis crassifolia* as described previously (Xu, et al., 2015). LG-134, a synthetic analogue of PCC, was also obtained from physachenolide D by chemical conversions (Zerio, et al., 2021). The purities of PCC and LG-134 were determined to be ≥ 98% by HPLC, and ^1^H and ^13^C NMR analyses.

### In vitro cytotoxicity assay

Serial two-fold dilutions of cells were plated into 96-well white-walled plates, and 72 hours later, the cells were incubated with CellTiter-Glo® 2.0 Cell Viability Assay (Promega, Madison, WI, Cat. No. G9242) reagent. There was a linear relationship (R^2^= 0.9647 to 0.9972) between the luminescent signal and the number of cells from 0 to 2000 cells/well. This linear range of the plot was used to determine the number of cells to plate for cytotoxicity assays. For cytotoxicity assays, cells were plated into 96-well white-walled plates and allowed to adhere for 24 hours before drug treatments. The cells were treated with serial dilutions of PCC or LG-134 in DMSO or vehicle control (DMSO) in presence or absence of poly I:C (10 µg/ml) and were incubated at 37LC for 48 hours. Cell viability was determined with the luminescence-based CellTiter-Glo 2.0 Assay. IC_50_ values were calculated as described in Statistics. To establish the limits of detection for ATP, two-fold serial dilution of ATP disodium salt (Sigma Cat.# A7699) in cell culture medium were added into 96-well white-walled plates and assayed as above. There was a linear relationship up to 10µM concentration, with an R^2^ estimated at 0.9982.

### In vivo treatment

To establish tumors, 1×10^6^ YUMM2.1 cells in 100 µl of PBS were injected intradermally (ID) into the right flank of 6 to 10 week-old male C57BL/6J or RAG-/-mice. Since YUMM2.1 cells are derived from male mice (Meeth, et al., 2016), only male mice were used to prevent recognition of Y-encoded antigens by the immune system of female mice. When tumor volume reached an average of >100 mm^3^ (6-9 days after tumor cell injection), mice were randomized into groups with equivalent average tumor volumes at treatment initiation. To determine the therapeutic efficacy of PCC in wild-type and RAG-/-mice, mice were treated with 20 mg/kg of PCC in 30% hydroxypropyl β-cyclodextrin (pharmaceutical grade Trappsol®, Cyclotherapeutics, Gainesville, FL), 30% DMSO and PBS (prepared as described – (Tewary, et al., 2021)) or vehicle control daily for 15 doses. For evaluation of the efficacy of PCC in combination with anti-PD-1 mAb, mice were treated with 1) vehicle control intratumoral (IT) daily for 15 doses and 10 µg/kg rat IgG2a, κ isotype control mAb (BioXCell, Lebanon, NH, Cat. No. BE0090, clone LTF-2) intraperitoneal (IP) every three days for the duration of the experiment; 2) PCC 20 mg/kg IT daily for 15 doses; 3) 10 µg/kg anti-PD-1 mAb (BioXCell, Cat. No. BE0146, clone RMP1-14) IP every three days for the duration of the experiment; or 4) PCC 20 mg/kg IT daily for 15 doses and 10 µg/kg anti-PD-1 IP every three days for the duration of the experiment.

The general health and tumor size and appearance were assessed daily, prior to the initiation of treatment; every other day, during PCC/vehicle control treatment; and thrice weekly after the completion of PCC/vehicle control treatment. Tumor volume was calculated using the formula volume = (L x W^2^)/2. Mice were weighed weekly, and n = 1 mouse (from the anti-PD-1 and PCC treatment group) was euthanized due to a weight loss of >20% in one week. Mice were euthanized when the volume of their primary tumor or recurrent tumor reached ≥ 2000 mm^3^.

### Flow cytometry

Cell apoptosis was detected by annexin V-fluorescein isothiocyanate (FITC) and 7-Aminoactinomycin D (7-AAD) double staining. YUMM2.1 cells were plated and were treated the following day with PCC (1.25 µM) or vehicle for 48 and 72 hours. The cells and the culture media were collected and stained with annexin V-FITC mAb (BioLegend, San Diego, CA, Cat. No. 640906) in binding buffer (10 mM HEPES, 140 mM NaCl, 2.5 mM CaCl_2_), followed by incubation with 7-AAD (BioLegend, San Diego, CA, Cat. No. 420403) in FACS buffer (1% bovine serum albumin and 0.05% sodium azide in PBS) prior to analysis.

For cell cycle analysis, cells were treated with PCC or vehicle for 48 hours as indicated in the cell apoptosis methods. Cells and their respective culture media were collected and were fixed and permeabilized with 70% ethanol at -20°C overnight, and resuspended in propidium iodide (PI)/RNAse (BD Biosciences, Franklin Lakes, NJ, Cat. No. 550825) prior to analysis. To determine the effect of drug washout, following 48 hours of PCC treatment, cells were washed and grown in media without PCC. The resulting cells and media were collected at 24 and 48 hours following washout and processed and stained as above. Results were analyzed using the Dean-Jett-Fox model for cell cycle data.

To assess the expression of PD-L1, YUMM2.1 cells were plated, and the next day grown in the presence or absence of 100 IU/mL recombinant murine IFN-γ (PeproTech, Cranbury, NJ, Cat. No. 315-05) for 24 hours. For dead cell exclusion, cells were stained with Fixable Viability Stain (FVS) 780 (BD Biosciences, Franklin Lakes, NJ, Cat. No. 565388) in PBS. FcγR III/II was blocked with anti-CD16/CD32 mAbs (BioLegend, San Diego, CA Cat. No. 101320) at 1 µg per 10^6^ cells in FACS buffer, stained with PD-L1-phycoerythrin (PE) mAb (eBiosciences, San Diego, CA, Cat. No. 12-5982-81), and placed in 1% paraformaldehyde prior to analysis.

For all flow cytometry experiments, cells were analyzed on a BD LSRII flow cytometer. Data were analyzed with BD FACS Diva and Flow Jo (BD Biosciences, Ashland, OR) software. Cells were first gated on FSC-A vs. SSC-A to remove debris, followed by stringent gating on FSC-A vs. FSC-H to remove doublets.

### TUNEL staining

To measure the double-stranded cleavage of DNA, the TUNEL assay was performed with the *In Situ* Cell Death Detection Kit, Fluorescein (Roche, Indianapolis, IN, Cat. No. 11684795910) following the manufacturer’s instructions. Briefly, YUMM2.1 cells were inoculated as described under *in vivo* treatment. When the tumor reached a volume of 100-550 mm^3^, mice received a single IT treatment of PCC (20 mg/kg) or vehicle. Twenty-four hours post treatment the mice were euthanized, and tumors were fixed in 10% neutral buffered formalin and embedded in paraffin. Following sectioning, deparaffinization and rehydration of tumor tissue, the tissue was microwave irradiated in 0.1 M citrate buffer, pH 6.0 for 5 minutes, cooled and rinsed twice in PBS. The tissues were then incubated with the TUNEL reaction mixture in a humidified chamber at 37°C for 1 hour. Subsequently, the tumor tissue was rinsed with PBS and mounted on microscope slides using ProLong Gold Antifade Mountant with DAPI (ThermoFisher) to stain for DNA and observed using a Zeiss AXIO Imager M2 epifluorescence microscope. For quantification of TUNEL staining, the percent of TUNEL positive/apoptotic cells was analyzed using Zen software. Approximately 10,000-25,000 DAPI positive cells were scored for TUNEL labeling.

### Statistics

Half maximal inhibitory concentration (IC_50_) values were calculated from the dose-response curve as the concentration of the drug that produced a 50% decrease in the mean luminescence relative to vehicle control wells. Curves were fit by nonlinear regression log-dose vs. normalized response with variable slope. The significance of differences between IC_50_ values were calculated using extra-sum-of-squares F test. Median time to onset of ulceration or tumor recurrence for each treatment group and mouse genotype was calculated using the Kaplan-Meier estimate, with comparisons using log rank tests. For the cell cycle and TUNEL experiments, the significance of differences between PCC vs. vehicle treated cells or tumors were calculated using t tests with Welch’s correction if n = 2 or the variances were not equal. These analyses were performed using GraphPad Prism software (San Diego, CA). Comparison of tumor volume and weight over time between treatment groups and mouse genotypes was assessed with a linear mixed effects model using Stata 15.0 (StataCorp, College Station, TX). A p value less than 0.05 was considered significant.

## Results

### PCC and LG-134 are cytotoxic to mouse melanoma cells

We tested whether immunogenic mouse melanoma cell lines YUMMER1.7, YUMMER.G and YUMM2.1, were sensitive to cell death induced by PCC (Figure 1A) and LG-134 (Figure 1B). Cell viability was determined based on quantitation of ATP, which is an indicator of metabolically active cells. Figure 1C shows the ATP standard curve, which demonstrates a linear luminescence signal for ATP concentrations from 0 to 10 µM. The linear range for cell number was determined (Figure 1D-F) and used to assess cytotoxicity. Cytotoxicity assays showed that YUMMER1.7 (IC_50_ = 0.5648 ± 0.1899 µM) (mean ± SD) and YUMM2.1 (IC_50_ = 0.5159 ± 0.1717 µM) were equally sensitive to PCC (p = 0.6857) (Figure 1G-I and Table 1). YUMMER.G (IC_50_ = 1.735 ± 0.1449 µM) was three-fold less sensitive to PCC treatment than YUMMER1.7 and YUMM2.1 (p = 0.0286). In addition to PCC, we tested whether the mouse melanoma cells were sensitive to its semi-synthetic analog, LG-134, which was found to have improved anti-tumor activity against prostate cancer cells (Zerio, et al., 2021). YUMM2.1 (IC_50_ = 0.1226 ± 0.0311 µM) had the highest sensitivity to LG-134 treatment, while YUMMER1.7 (IC_50_ = 0.2852 ± 0.0680 µM) were slightly less sensitive to LG-134 mediated cell death. As with PCC treatment, YUMMER.G cells (IC_50_= 0.4500 ± 0.1996 µM) had the least sensitivity to LG-134-induced cell death. Treatment with PCC has previously been shown to amplify TRAIL-mediated caspase-8-dependent extrinsic apoptosis signaling in human renal carcinoma cells (Xu, et al., 2017). A number of reports have suggested that the TLR3 ligand poly I:C can promote caspase-8-dependent apoptosis in certain human cancers such as melanomas and breast carcinomas (Estornes, et al., 2012, Salaun, et al., 2007, Tewary, et al., 2021, Weber, et al., 2010). We, therefore, tested whether the addition of poly I:C can enhance the sensitivity of YUMMER1.7, YUMMER.G or YUMM2.1 melanoma cells to PCC and LG-134. The addition of poly I:C did not enhance the sensitivity of these cells to PCC-induced cell death; however, the addition of poly I:C slightly enhanced the sensitivity of these cells to LG-134-induced cell death by 1.5-fold (Figure S1 and Table 1). In comparing the effects of PCC vs. LG-134 treatment, all three cell lines were significantly more sensitive to LG-134 than PCC (Table 1). YUMMER.G and YUMM2.1 cells displayed the highest increase in sensitivity to LG-134 (2.63-2.75 – fold higher in LG-134 than PCC), and YUMMER1.7 cells were only slightly more sensitive (1.43-fold higher in LG-134 than PCC). These data demonstrate that both PCC and LG-134 are effective cytotoxic agents with the potential to treat melanoma tumors *in vivo*.

**Figure 1.**
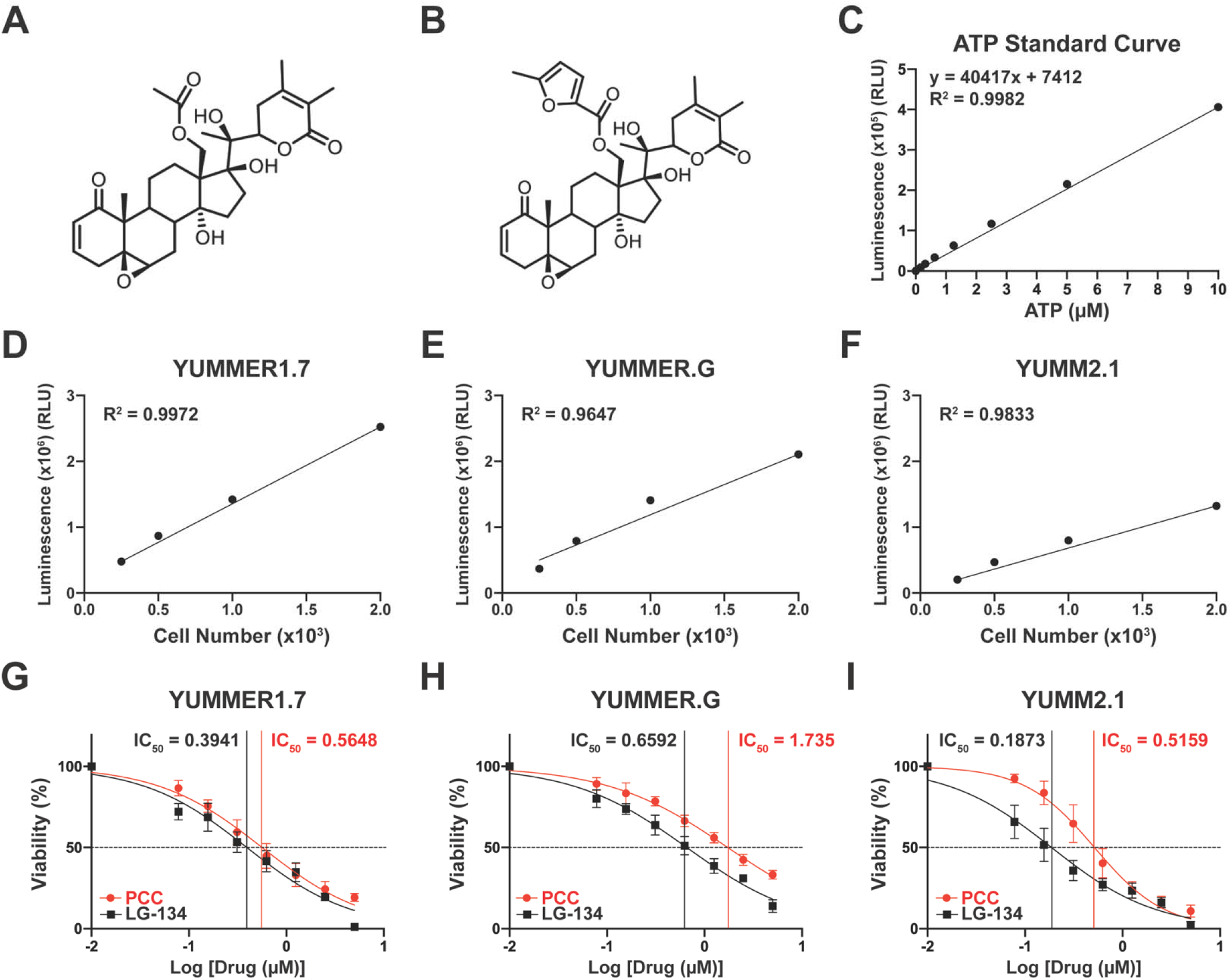
Mouse melanoma cells are sensitive to PCC and LG-134 induced death. Chemical structures of physachenolide C (PCC) (A) and LG-134 (B). (C) ATP standard curve using CellTiter-Glo assay. (D-F) Cell number titration of mouse melanoma cell lines. (G-I) Mouse melanoma cell lines were treated with increasing concentrations of PCC or LG-134 for IC_50_ determination. Cell viability was measured after 48 hours of treatment. Graphs show the percent viability (mean ± SD) of four independent experiments for each cell line. IC_50_ values were calculated from the mean of four independent experiments for each cell line using non-linear fit of log-dose vs. normalized response with variable slope.

**Table 1.**
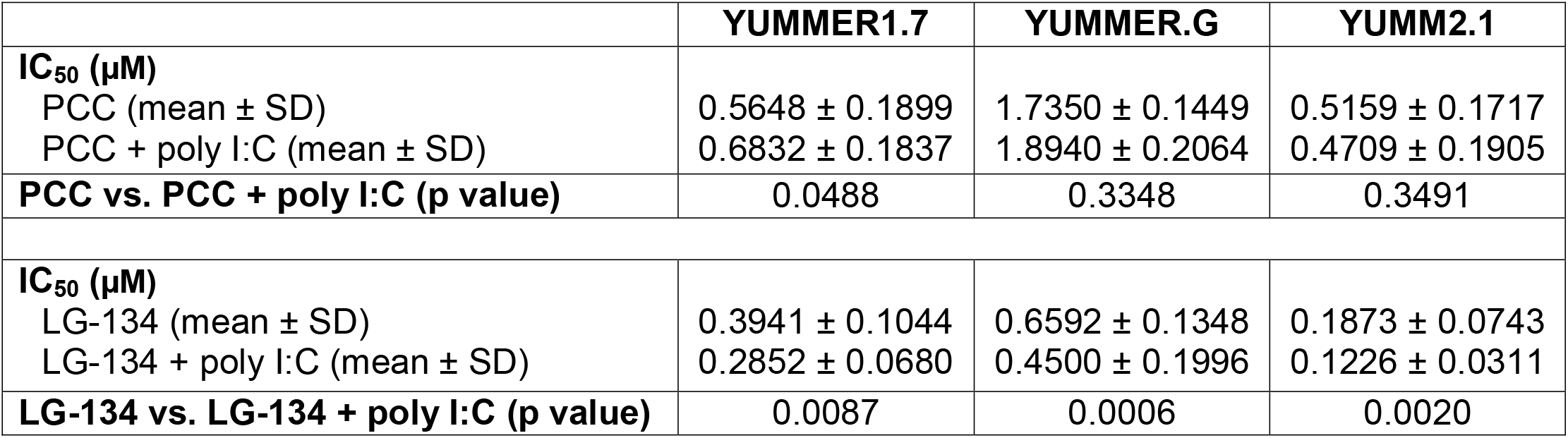
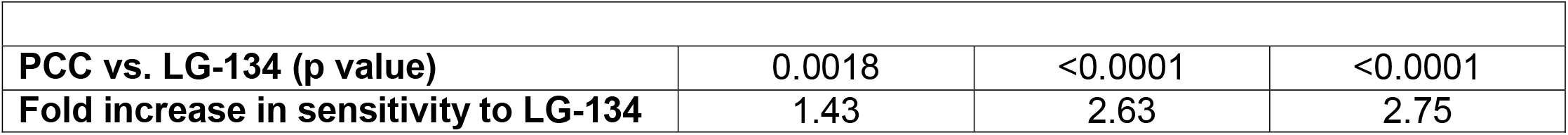
Half maximal inhibitory concentration (IC_50_) for PCC and LG-134 in murine melanoma cells

|  | YUMMER1.7 | YUMMER.G | YUMM2.1 |
| --- | --- | --- | --- |
| **IC<sub>50</sub> (μM)** |  |  |  |
| PCC (mean ± SD) | 0.5648 ± 0.1899 | 1.7350 ± 0.1449 | 0.5159 ± 0.1717 |
| PCC + poly I:C (mean ± SD) | 0.6832 ± 0.1837 | 1.8940 ± 0.2064 | 0.4709 ± 0.1905 |
| **PCC vs. PCC + poly I:C (p value)** | 0.0488 | 0.3348 | 0.3491 |
| **IC<sub>50</sub> (μM)** |  |  |  |
| LG-134 (mean ± SD) | 0.3941 ± 0.1044 | 0.6592 ± 0.1348 | 0.1873 ± 0.0743 |
| LG-134 + poly I:C (mean ± SD) | 0.2852 ± 0.0680 | 0.4500 ± 0.1996 | 0.1226 ± 0.0311 |
| **LG-134 vs. LG-134 + poly I:C (p value)** | 0.0087 | 0.0006 | 0.0020 |
| **PCC vs. LG-134 (p value)** | 0.0018 | <0.0001 | <0.0001 |
| **Fold increase in sensitivity to LG-134** | 1.43 | 2.63 | 2.75 |

**Supplemental Figure 1.**
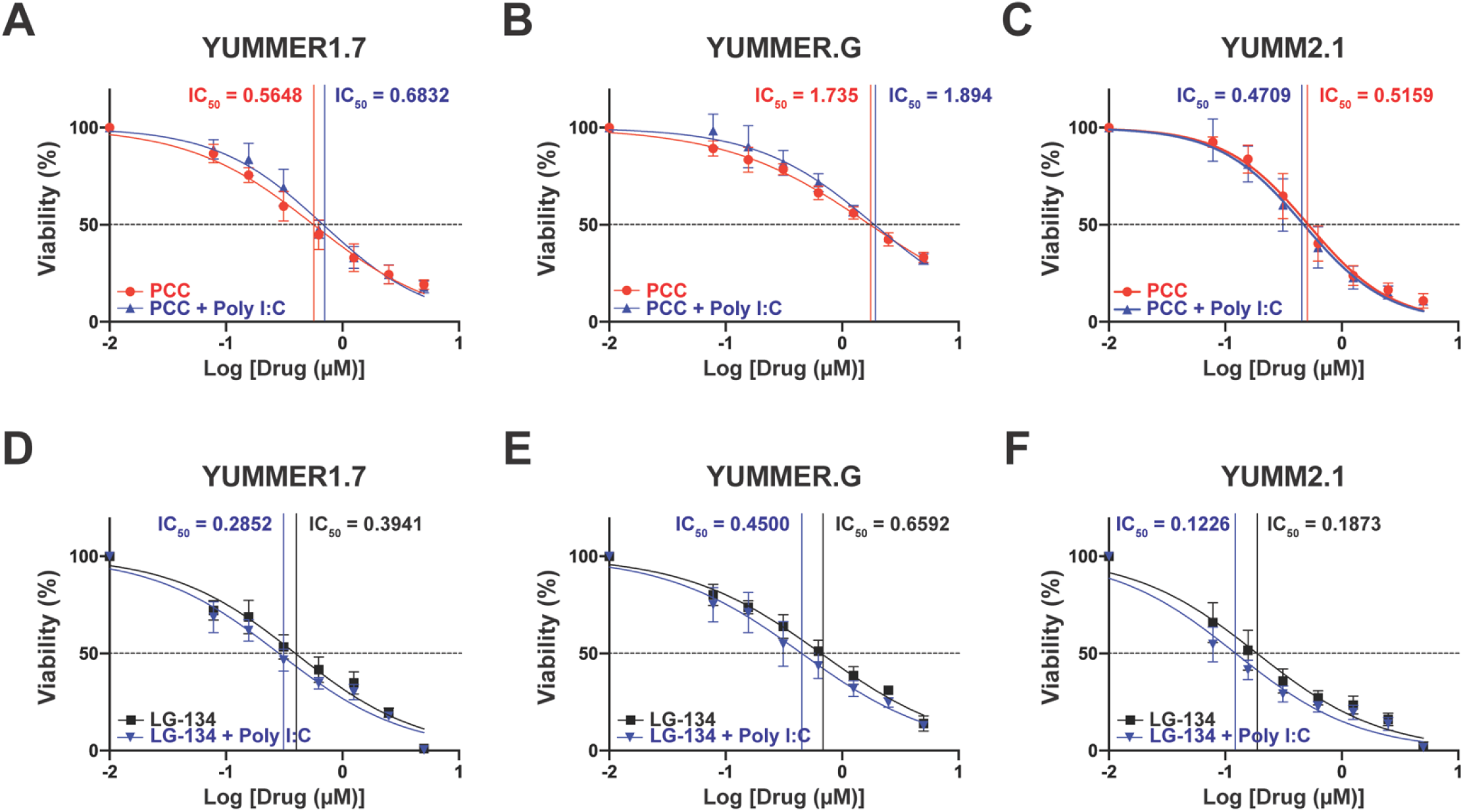
Cytotoxicity assay of mouse melanoma cell lines treated with PCC or LG-134 alone or with poly I:C. Mouse melanoma cell lines were treated with increasing concentrations of PCC or LG-134 alone or in combination with poly I:C (10 µg/ml). Cell viability was measured after 48 hours of treatment. Graphs show the percent viability (mean ± SD) of four independent experiments for each cell line. IC_50_ values were calculated from the mean of four independent experiments for each cell line using nonlinear fit of log-dose vs. normalized response with variable slope.

### PCC treatment causes complete regression of established melanoma tumors

We selected to proceed with testing PCC in the treatment of tumors *in vivo*, given that PCC is easier to obtain, has improved solubility (data not shown), and is less toxic to normal cells than LG-134 (Zerio, et al., 2021). We selected the YUMM2.1 melanoma model, over YUMMER1.7 and YUMMER.G, for *in vivo* testing, given that YUMM2.1 cells exhibit more consistent *in vivo* tumor formation. Injection of 1 x 10^6^ YUMM2.1, YUMMER1.7 and YUMMER.G cells formed progressively, enlarging tumors in 17/19 (89%), 7/10 (70%), and 11/15 (73%) mice, respectively (data not shown). Based on a prior *in vivo* study with prostate cancer xenografts using five times per week dosing of PCC, which also showed that the serum half-life of PCC after subcutaneous administration is 2 to 3 hours (Xu, et al., 2015), we opted to treat established YUMM2.1 tumors IT daily. Following ID injection of 1×10^6^ YUMM2.1 cells and when tumors reached an average volume of greater than 100 mm^3^, treatment with PCC at 20 mg/kg IT daily resulted in the complete regression of clinically apparent tumors in 100% of wild-type mice (Figure 2A). Over three experiments, all 15 wild-type mice treated with PCC had complete regression of tumors within 10.1 ± 0.5 days (mean ± SEM). Substantial toxicity to PCC was not observed, as the PCC treated mice appeared in general good health and did not experience weight loss (Figure 2B) or diarrhea (data not shown). In each of three experiments, the weight of PCC treated mice over time relative to the weight at tumor injection was equal to or greater than the weight of vehicle treated mice. The YUMM2.1 melanoma model was prone to ulceration (Figure 2C). Ulceration at the site of the tumor/injections was observed in all mice, regardless of treatment group. However, the median onset of ulceration in the PCC treated group was earlier than in the vehicle control group, suggesting that the tumor response to PCC or PCC itself may lead to ulceration. These data demonstrate the *in vivo* efficacy of PCC in generating a complete response in the YUMM2.1 murine melanoma model.

**Figure 2.**
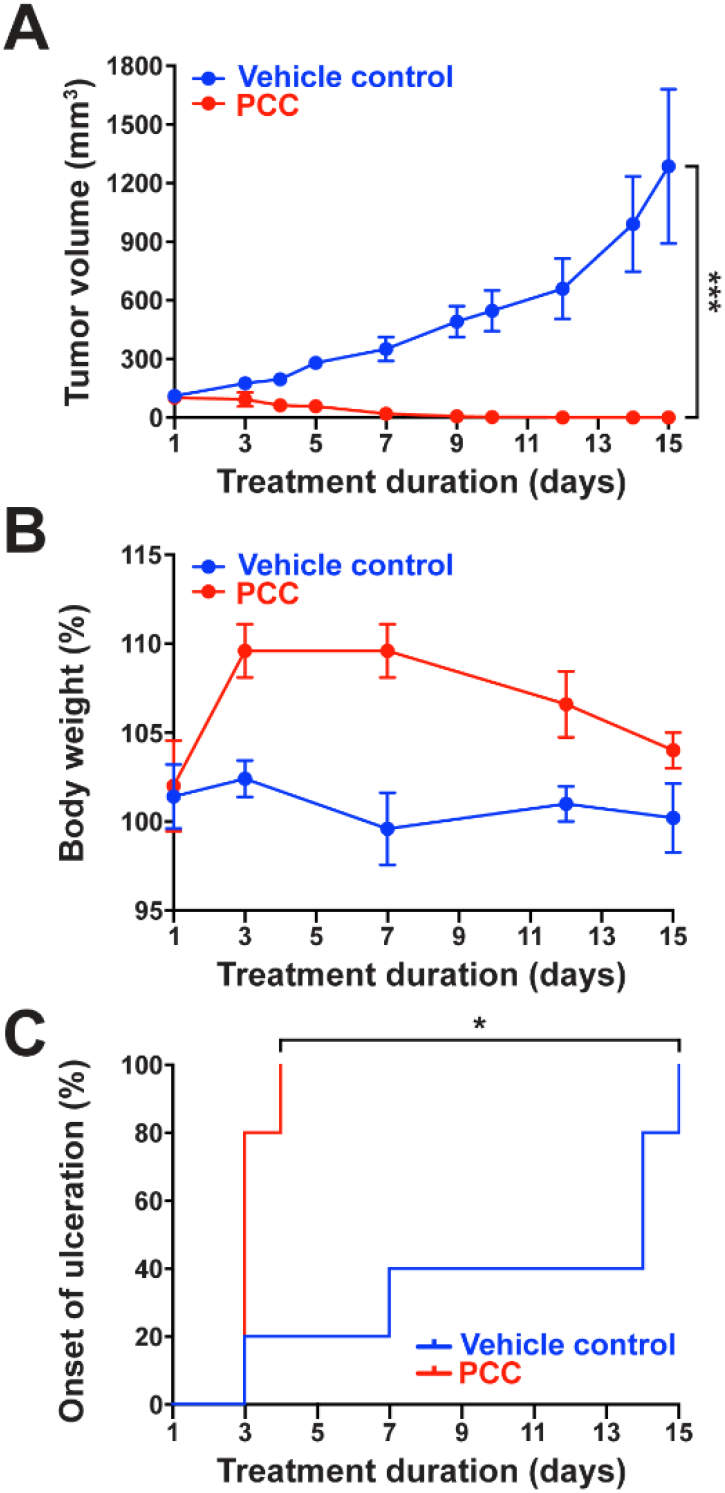
Melanoma tumors completely regress after treatment with PCC. (A) Tumor volume (mean ± SEM) for mice with established YUMM2.1 melanoma tumors treated with either PCC 20 mg/kg or vehicle control IT daily from day 1 through day 15. (p < 0.001; linear mixed effects model) (B) Body weight, measured as a percentage of the weight at the time of tumor cell injection, was followed throughout treatment. (C) Onset of ulceration at the tumor/site of injection (p = 0.013; log rank). Data shown are from one experiment and representative of three experiments each with n = 5 mice per group. *, p < 0.05; ***, p < 0.001

### PCC treatment induces apoptosis in vitro and in vivo

To evaluate the mechanism of PCC-induced cell death, we assessed the ability of PCC to induce apoptosis. First, we determined the percentage of YUMM2.1 cells undergoing apoptosis *in vitro* following PCC treatment, using flow cytometric analysis of annexin V/7-AAD staining of cells. PCC treatment resulted in a 4.7-fold increase in the percentage of early apoptotic cells at 48 hours and a 10-fold increase at 72 hours compared to vehicle treated cells (Figure 3A-B). Similarly, there was a 1.7-fold increase in late apoptotic cells in PCC treated cells compared with vehicle treated cells at 48 hours, and 3.3-fold increase at 72 hours. These data demonstrate that PCC treatment induces apoptosis *in vitro*, and that proportionally more cells are undergoing apoptosis at 72 hours compared to 48 hours when normalized to vehicle control. Next, we quantified the level of apoptosis *in vivo* 24 hours after YUMM2.1 tumors were treated with PCC (20 mg/kg) or vehicle control IT, using the TUNEL assay which detects breaks in DNA strands during late apoptosis. Within the tumor, DAPI positive cells were quantified, and the percentage of cells that were TUNEL-positive was determined. In tumors that were treated with vehicle alone, a very small fraction (mean of 0.8%) of cells were TUNEL-positive. In contrast, tumors treated with PCC had a significantly increased percentage of cells (40%) that were TUNEL-positive (Figure 3C-D). Taken together, these data demonstrate that PCC induces apoptosis in melanoma cells *in vitro* and *in vivo*.

**Figure 3.**
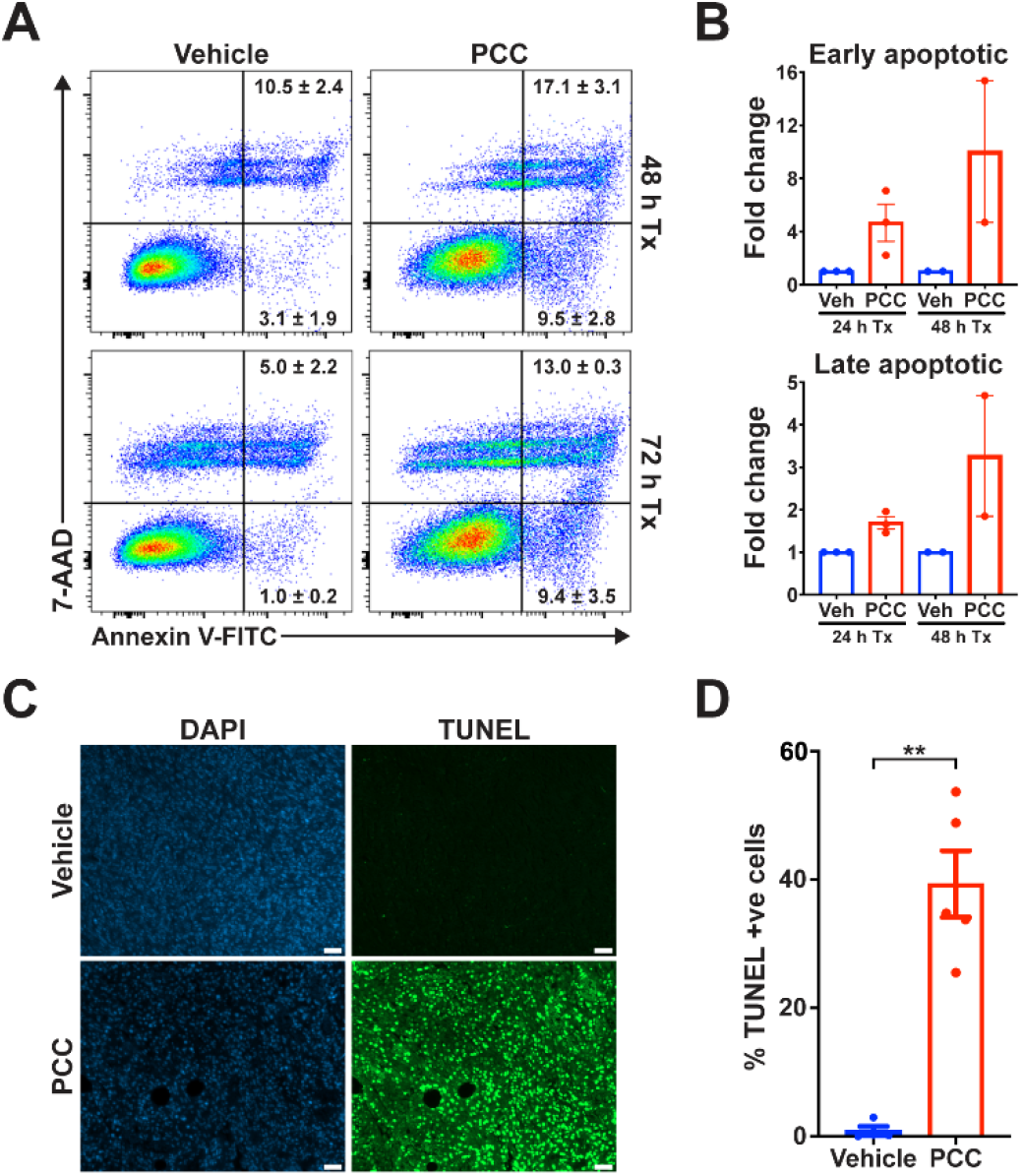
PCC treatment induces apoptosis in YUMM2.1 cells *in vitro* and *in vivo*. (A) Representative flow cytometry plots showing staining with apoptotic marker annexin V and DNA stain 7-AAD. Cells were grown with and without 1.25 µM PCC treatment (Tx) for 48 and 72 hours prior to analysis. Early apoptotic cells are defined as annexin V+/7-AAD- and late apoptotic cells are defined as annexin V+/7-AAD+. Numbers indicate the mean ± SEM from 2-3 experiments. (B) Fold change of early and late apoptotic cells in PCC treated cells normalized to vehicle treated control cells. Bar graphs show the mean ± SEM; points show the values from each independent experiment. (C) Representative images of DAPI and TUNEL staining in formalin-fixed, paraffin-embedded YUMM2.1 tumors 24 hours after PCC (20 mg/kg) or vehicle control IT treatment. Scale bar, 100 µm (D) Quantitation of the TUNEL-positive apoptotic cells out of the DAPI-positive cells within the tumor. Significance was assessed by a t test using Welch’s correction (p = 0.0016). Bar graph shows the mean ± SEM; each point represents the value from single tumor. **, p < 0.01

### T cell-mediated immunity does not contribute to the therapeutic efficacy of PCC or prevent tumor recurrence

All mice treated with PCC had complete regression of the clinically apparent tumor, and the response was durable in a portion of the mice after discontinuation of treatment (Figure 4A). Over three experiments with a total of 15 wild-type mice treated with PCC, the tumor recurred in 67% of mice at 13.7 ± 1.9 days (mean ± SEM) after discontinuing treatment. To assess the immunogenicity of the YUMM2.1 model, we evaluated *in vivo* tumor growth in wild-type mice compared to RAG-/-mice, lacking T and B cells. Following injection of a small number of YUMM2.1 cells (1 x 10^5^), there was increased tumor growth in RAG-/-mice, compared to wild-type mice, demonstrating that the adaptive immune response can constrain *in vivo* YUMM2.1 tumor growth (p < 0.001; linear mixed effects model) (Figure 4B). Next, we evaluated whether the adaptive immune response contributed to the therapeutic efficacy of PCC or delayed the tumor recurrence following PCC treatment. Wild-type and RAG-/-mice with established YUMM2.1 tumors were treated with a 15 day course of PCC. The decrease in tumor volume during treatment was not improved in wild-type, compared to RAG-/-, mice, and there was no difference in the increase in tumor volume after discontinuation of PCC treatment between wild-type and RAG-/-mice (Figure 4C). Additionally, there was no difference in the recurrence free survival following PCC treatment between wild-type and RAG-/-mice (p = 0.71; log rank; data not shown). These results demonstrate that the adaptive immune response does not contribute to the therapeutic efficacy of PCC or prevent recurrence in the YUMM2.1 model of melanoma. To further investigate whether the immune response contributed to the durable responses to PCC observed in 33% of mice, mice with a complete response to PCC were rechallenged with YUMM2.1 cells to test for immunologic memory, a feature of the adaptive immune response (Figure 4D). YUMM2.1 tumors grew in all mice after re-challenge, but tumor growth in the rechallenged mice was delayed compared to naïve mice (p < 0.001; linear mixed effects model). These results may suggest partial immune memory. We hypothesized that the combination of PCC with an anti-PD-1 mAb, to increase T cell responses, might diminish tumor recurrence. YUMM2.1 cells expressed PD-L1, the ligand for PD-1, following exposure to IFN-γ *in vitro* (Figure 4E). However, the combination of PCC with an anti-PD-1 mAb did not improve the therapeutic efficacy (p = 0.10; linear mixed effects model), did not diminish recurrence after discontinuation of PCC (p = 0.70; linear mixed effects model), and did not improve the recurrence free survival (p = 0.75; log rank), compared with PCC alone in the YUMM2.1 model of melanoma (Figure 4F). While a partial response of YUMM2.1 tumors to anti-PD-1 mAb treatment has been previously reported (Homet Moreno, et al., 2016), we did not observe any difference in tumor growth between the anti-PD-1 mAb alone vs. the vehicle/isotype control groups (p = 0.20; linear mixed effects model). Together, these results show that T cell-mediated immunity does not contribute to the therapeutic efficacy of PCC in YUMM2.1 model.

**Figure 4.**
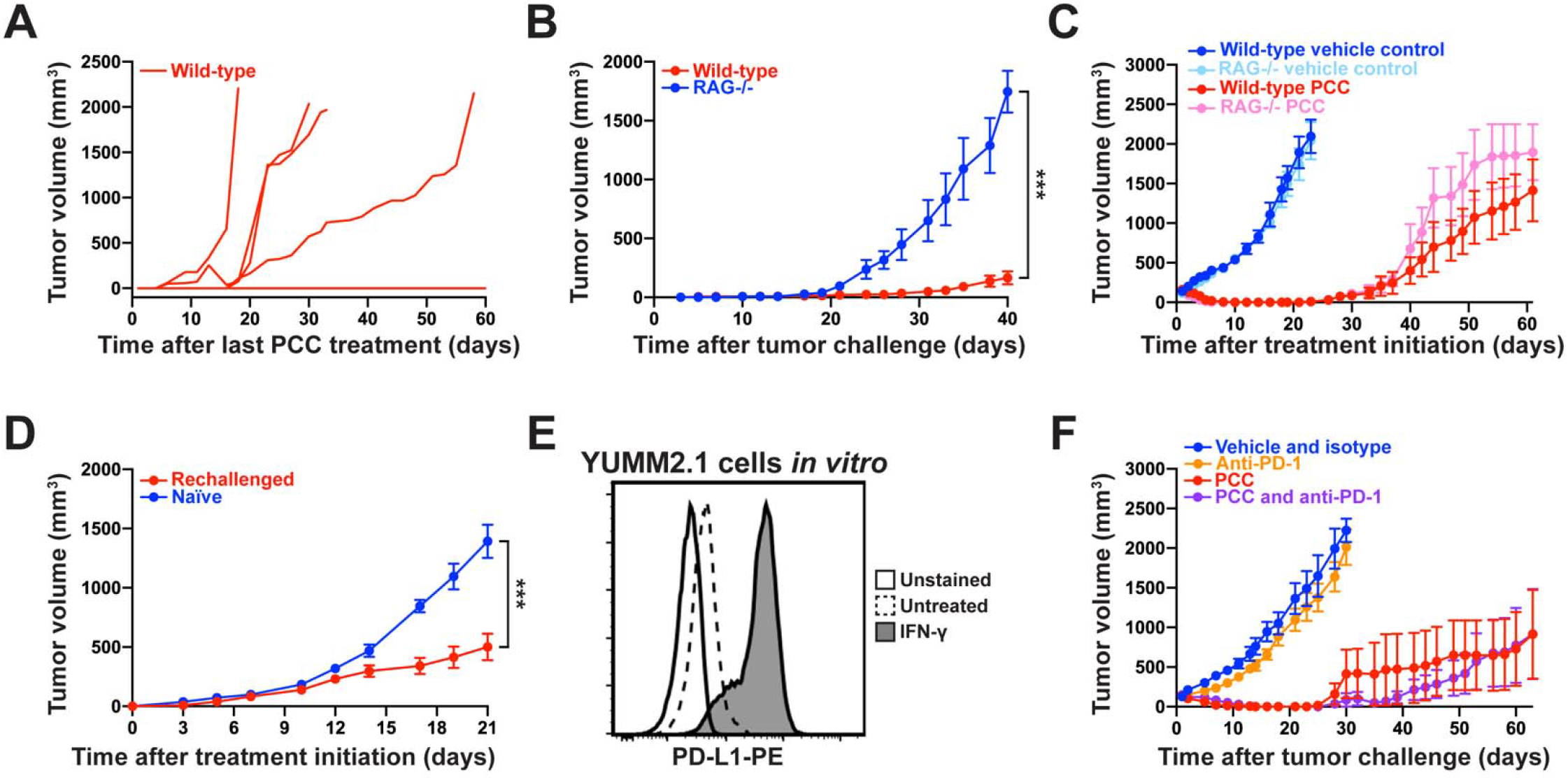
T cell-mediated immunity does not contribute to the therapeutic efficacy of PCC or prevent tumor recurrence. (A) Long term follow-up after discontinuation of PCC treatment. Mice with established YUMM2.1 tumors were treated with a 15 day course of daily PCC IT as in Figure 2A. Graph shows the individual tumor growth curves where day 1 is the first day off treatment. Data shown are from one experiment and representative of three experiments, each with n = 5 mice per group. (B) Tumor growth following ID injection of 1 x 10^5^ YUMM2.1 cells in wild-type vs. RAG-/-mice. Data shown as tumor volume (mean ± SEM) from one experiment with n = 5 mice per group. (C) Effect of PCC treatment on melanoma tumors in wild-type vs. RAG-/-mice. Following injection of 1 x10^6^ YUMM2.1 cells, mice received daily PCC IT treatment at 20 mg/kg from day 1 through day 15. Graph shows tumor volume (mean ± SEM) over time, using last observation carried forward to produce smoothed curves. Data shown are from one experiment and representative of two experiments with n = 5 mice per group in each. (D) Tumor volume (mean ± SEM) over time in rechallenged vs. naïve mice. Mice that remained free of tumor for 59 -104 days after completion of a 15 day course of PCC treatment and naïve mice received ID injection of 1 x10^6^ YUMM2.1 cells. Data shown from one experiment with n = 6 mice per group. (E) PD-L1 expression is IFN-γ-inducible (100 IU/mL) following 24 hours of treatment on YUMM2.1 cells *in vitro* by flow cytometry. Data shown are representative of four experiments. (F) Comparison of tumor growth in mice with YUMM2.1 tumors treated with combination of PCC and anti-PD-1 in comparison to PCC alone, anti-PD-1 alone or vehicle/isotype control. Mice treated with PCC received daily treatment from day 1 through day 15. Mice treated with anti-PD-1 received 10 µg/kg IP every three days, starting on day 1 for the duration of the experiment. Graph shows tumor volume (mean ± SEM) over time, using last observation carried forward. Data shown are from one experiment with n = 5 mice per group. ***, p < 0.001

### PCC causes arrest of melanoma cells in the G0-G1 phase

Our findings above show that while PCC treatment causes complete tumor regression, some tumors eventually recur after cessation of treatment in a T-cell independent manner. We, therefore, hypothesized that PCC treatment could cause a concomitant increase in cell death and growth-arrest of the surviving population, which can re-enter cell cycle upon removal of the treatment. To test the effect of PCC on the cell cycle of melanoma cells, YUMM2.1 cells were treated *in vitro* with PCC for 48 hours and followed for an additional 48 hours post-wash-out of the drug. Consistent with Figure 1 and 3, YUMM2.1 cells treated with PCC for 48 hours showed an increase in cell death (Figure 5A). Although there was an increase in cell death in the presence of PCC, there was also a concomitant increase in G0-G1 arrested cells. In the PCC treated cells, a higher fraction of cells accumulated in the G0-G1 phase compared to vehicle (52.9 ± 3.5% (mean ± SEM) in PCC treated vs. 27.0 ± 1.5% vehicle, p = 0.0024) (Figure 5B-C). Furthermore, this increase in G0-G1 fraction was accompanied by a decrease in the percentage of cells in S phase (19.5 ± 1.9% in PCC treated vs. 39.3 ± 2.2% vehicle, p = 0.0023), and a smaller decrease in the percentage of cells in the G2-M phase compared to vehicle treated cells implying an exit from cell cycle. These findings suggest that while PCC treatment induces apoptosis in a fraction of the melanoma cells, it also drives the remaining cells into a G0-G1 arrest. However, this growth arrest was only transient. Following washout of the drug, PCC treated cells were able to re-enter cell cycle and proliferate similar to vehicle treated cells as early as 24 hours post wash-out (Figure 5A-C), as evidenced by a decrease in the percentage of cells in G0-G1 in post-PCC treated samples to levels comparable to vehicle treated cells at 24 hours (41.2 ± 2.5% PCC treated vs. 32.1 ± 3.6% vehicle, p = 0.1839). Additionally, there was a significant increase in the percentage of cells entering S-phase in PCC treated cells at 24 hours post-washout, such that there was no longer a difference in the percentage of cells in S-phase between PCC treated (29.4 ± 1.4%) vs. vehicle treated (35.5 ± 1.1%) cells (p = 0.083). Furthermore, by 48 hours following PCC removal, the cells had cell cycle profiles almost identical to the vehicle treated cells that were actively proliferating (G0-G1 33.8 ± 5.0% PCC treated vs. 33.5 ± 0.7% vehicle, p = 0.9680 and S phase 34.3 ± 3.5% PCC treated vs 32.7 ± 1.9% vehicle, p = 0.74). These data demonstrate that PCC treatment induces G0-G1 cell cycle arrest in YUMM2.1 cells and that the cell cycle progression is normalized after PCC is removed.

**Figure 5.**
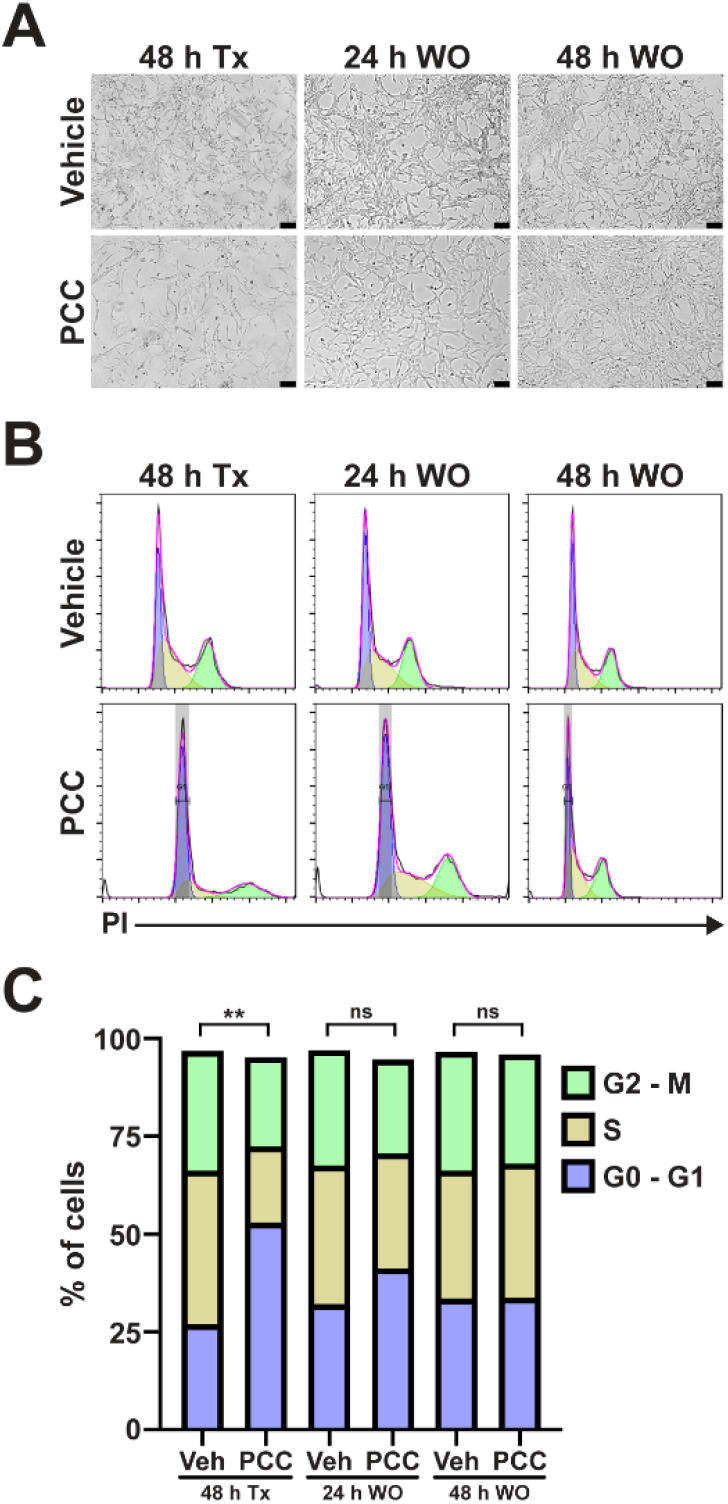
PCC causes G0-G1 arrest in YUMM2.1 melanoma cells. (A) Representative pictures of cells grown in culture with and without PCC treatment (1.25 µM) at 48 hours (Tx), and at 24 and 48 hours after washout (WO). Scale bar, 200 µm (B) Representative PI staining and cell cycle models showing the percentage of cells in G0-G1 (purple), S (tan), and G2-M (green) phases of the cell cycle. (C) Percentage of cells in G0-G1, S and G2-M following 48 hours of PCC treatment and at 24 and 48 hours after WO. The percentage of cells in G0-G1 were compared between PCC and vehicle (Veh) treated cells for 48 hours after treatment and at 24 and 48 hours after WO. Data are shown from two to three experiments per condition. **, p < 0.01; ns, not significant

## Discussion

Using an immunogenic mouse melanoma model, we show that the natural product PCC is a potent suppressor of melanoma tumor growth. PCC suppresses melanoma tumor growth by concomitantly activating apoptotic and growth suppressive programs in the tumor cells. PCC treatment of established melanoma tumors results in complete regression of tumors in all mice, with a durable response in 33% of mice.

Previous findings have shown that twice weekly treatment with PCC alone for three weeks resulted in only partial regression of melanoma tumors when using a human melanoma xenograft model and a non-immunogenic mouse melanoma cell line (Tewary, et al., 2021). However, PCC mediated cell death and tumor regression in these models were significantly enhanced by the addition of an immune stimulant poly I:C. Herein we demonstrate that treatment with PCC alone daily for 15 days causes complete regression of melanoma tumors in a syngeneic mouse melanoma model. Our findings indicate that PCC can be an effective therapy against melanoma. We observe a significant increase in apoptosis of the melanoma cells both *in vitro* and *in vivo* as evidenced by an increase in annexin V-positive and TUNEL-positive cells. Previous studies examining the mechanisms of PCC-induced apoptosis demonstrated that PCC sensitizes melanoma and other cancer cells to apoptosis by enhancing caspase-8-dependent apoptosis signaling and reducing the levels of anti-apoptotic proteins cFLIP (also known as CLFAR (CASP8 and FADD Like Apoptosis Regulator)) and BIRC7 (Baculoviral IAP repeat containing 7, also known as Livin or ML-IAP (Melanoma-Inhibitor of Apoptosis)) (Tewary, et al., 2021). Both cFLIP and Livin have been shown to be regulated by bromo and extraterminal domain (BET) protein BRD4 and BET inhibitor JQ1, and, thereby, enhance resistance to therapy in several cancers (Sugihara, et al., 2020, Yao, et al., 2015). PCC was shown to bind and inhibit the biological activity of the BET proteins, specifically BRD4 protein, thereby leading to a decrease in levels of anti-apoptotic inhibitors of apoptosis proteins (IAPs), such as Livin and c-FLIP, and resulting in increased tumor cell death and tumor regression (Tewary, et al., 2021). However, upon discontinuation of PCC, YUMM2.1 cells and tumors were able to proliferate and recur both *in vitro* and *in vivo*. The lack of PCC regulated apoptosis signals could contribute towards the increased survival of these cells and subsequent proliferation. But, other apoptosis independent mechanisms are likely to have a role in this process. We found that PCC treatment resulted in G0-G1 cell cycle arrest. In addition to regulating apoptosis signaling, BRD4 has also been shown to regulate genes involved in cell cycle and cell proliferation. Cyclin D1, a major regulator of cell cycle G1-S transition, and CDC25A, a phosphatase required for G1-S transition are upregulated by BRD4 in cutaneous squamous cell carcinoma (Xiang, et al., 2018) and hepatocellular carcinoma (Hong, et al., 2016). We, therefore, postulate that PCC mediated suppression of BRD4 can also lead to decreased activity of Cyclin D1 and CDC25A leading to arrest in G0-G1. The recovery of normal activity of these cell cycle regulators following drug wash-out may play a role in the proliferation and relapse of melanoma tumors *in vivo*. Therefore, the role of these cell cycle mediators in tumor growth suppression and subsequent recurrences must be further investigated. The addition of the immune stimulant poly I:C was shown to enhance the apoptotic effects of PCC in human melanoma cells both *in vitro* and *in vivo* and lead to complete tumor regression for up to 3 months (Tewary, et al., 2021). While we did not see any additional effect of adding poly I:C on PCC mediated cell death *in vitro*, it is possible that poly I:C mediated immune stimulation *in vivo* could prolong the effects of PCC and suppress recurrences.

In the YUMM2.1 model, T cell-mediated immunity did not contribute to the therapeutic efficacy of PCC or diminish tumor recurrence. The lack of T cells in RAG-/-mice did not diminish the efficacy of PCC or reduce tumor recurrence, compared to wild-type mice. Additionally, the combination of PCC with an anti-PD1 mAb did not enhance the efficacy or reduce tumor recurrence, compared with PCC alone. Previous work showed that PCC sensitizes B16 murine melanoma cells to cell death in response to the combination of TNF-α and IFN-γ and that PCC sensitizes human melanoma cell lines to cell death from soluble factors produced by activated T cells, including TNF-α (Tewary, et al., 2021). Based on these findings, the authors of the prior study hypothesized that PCC may augment the efficacy of T cell-mediated immunotherapies, which we did not observe. One explanation is that T cells are not necessary as the source of TNF-α and IFN-γ *in vivo*. Activated macrophages and dendritic cells are major producers of TNF-α, and other cell types also produce this cytokine in tumors, including tumor cells and fibroblasts (www.immgen.org) (Heng and Painter, 2008, Wajant, 2009). Natural killer cells are another a major source of IFN-γ within tumors (www.immgen.org) (Alspach, et al., 2019). In addition to T cells, whether other immune cells, such as macrophages, dendritic cells, and natural killer cells, could have a role in enhancing the therapeutic efficacy of PCC remains to be investigated. Alternatively, the melanoma model in this study, which is distinct from the models previously tested, may be less sensitive to cytokine-induced cell death. Consistent with the idea that different models have different susceptibility to cell death pathways, the addition of poly I:C significantly enhanced the cytotoxicity of PCC in human melanoma cell lines *in vitro* and the efficacy of PCC in treating B16 *in vivo* (Tewary, et al., 2021), whereas poly I:C did not enhance the sensitivity to PCC in any of the three melanoma models that we tested. The YUMM2.1 model is immunogenic, in that we showed that tumor growth is reduced in RAG-/-mice, compared to wild-type mice, and that tumor growth is delayed in rechallenged mice compared to naïve mice. If anti-tumor T cell responses contribute to improved tumor control with PCC, selection of an alternative melanoma model with a greater tumor mutational burden might better allow the detection a synergistic effect of PCC and T cell-mediated immunotherapy.

This study demonstrates that continuous treatment with PCC can effectively treat established melanoma tumors and suppress melanoma tumor growth with minimal toxicity. Thus, PCC and other modifications of 17β-hydroxywithanolides warrant further development to improve the therapeutic outcome for patients with advanced melanoma.

## Acknowledgments

Flow cytometry was performed at the Flow Cytometry Core directed by Dr. Mrinalini Kala at the University of Arizona College of Medicine Phoenix. This work was supported by the Arizona Biomedical Research Commission (ADHS-16-162515) and the National Institutes of Health (P30-CA023074).

## Abbreviations

BET: bromo and extraterminal domain
FITC: fluorescein isothiocyanate
IAP: anti-apoptotic inhibitors of apoptosis proteins
IFN-γ: interferon gamma
ID: intradermal
IP: intraperitoneal
IT: intratumoral
mAb: monoclonal antibody
PCC: physachenolide C
PE: phycoerythrin
PI: propidium iodide
YUMM: Yale University Mouse Melanoma
7-AAD: 7-Aminoactinomycin D

